# Bacterial metagenome in plaque, saliva, and tumor samples from individuals with and without OSCC by next-generation sequencing

**DOI:** 10.64898/2026.08.27.747557

**Authors:** Alveiro Erira, Dabeiba Adriana García Robayo, Fredy Gamboa, Andrés Ignacio Chala, Andrei Moreno, Angel Cid Arregui, Eliana Muñoz, José Noguera, Fabian Tobar Tosse

## Abstract

**Background:** Oral dysbiosis has been associated with oral squamous cell carcinoma (OSCC); however, most microbiome studies rely on 16S ribosomal RNA (rRNA) gene sequencing, limiting species-level taxonomic resolution.

**Methods:** Dental plaque, saliva, and tumor tissue samples from 10 patients with OSCC and dental plaque and saliva samples from 10 healthy controls were analyzed in this exploratory cross-sectional study. DNA was extracted and subjected to shotgun metagenomic sequencing using the Illumina MiSeq platform. Sequence reads were quality filtered with fastp, taxonomically classified using Kraken2 v2.1.3, and species-level abundances were re-estimated with Bracken v2.9 following the removal of human reads and low abundance taxa. Relative abundances were compared using the Mann–Whitney U test with the Benjamini– Hochberg false discovery rate correction, while the Bray–Curtis principal coordinate analysis was used as an exploratory approach to visualize microbial community patterns.

**Results:** Shotgun metagenomic sequencing revealed distinct bacterial community profiles across the oral microenvironment. Dental plaque exhibited the highest taxonomic diversity and relative abundance. The control plaque was enriched in *Streptococcus koreensis, Capnocytophaga sp*. oral taxon 878, *Treponema sp*. *Marseille* Q4132, and *Leptotrichia sp.* oral taxon 498, whereas the plaque from patients with OSCC showed a higher relative abundance of *Pyramidobacter piscolens*, *Parvimonas parva*, and *Gemella sanguinis*. Salivary samples displayed lower diversity and a more homogeneous composition, predominantly comprising *Capnocytophaga endodontalis*, *Prevotella jejuni*, *Aggregatibacter aphrophilus*, and *Gemella sanguinis*. The tumor tissue showed relatively higher abundance of *Sellimonas catena*, *Escherichia coli, Solobacterium moorei*, and *Lacrimispora sp*. HJ-01.

**Conclusions:** This exploratory study provides species-level characterization of the oral microbiome across multiple oral microenvironments in OSCC and generates hypotheses for future integrative metagenomic and functional studies investigating the potential contribution of oral bacterial communities to OSCC pathogenesis.

## CONTEXT

Oral squamous cell carcinoma (OSCC) is the most common oral cancer [1,2] and is associated with chemical, physical, and biological risk factors [2–5]. Among these, the oral microbiome stands out for its ability to induce chronic inflammation, mutagenesis, and oncogenic activation, as well as promoting angiogenesis and cell proliferation [4–7]. Oral eubiosis is associated with health, while dysbiosis can contribute to the development of diseases, including cancer [8,9]. Various studies [4,10–12] have shown that an imbalance in the oral microbiome can increase the risk of developing cancer by inducing uncontrolled cell proliferation, and cytoskeletal alterations, inhibiting apoptosis, and promoting mutagenesis. Additionally, bacterial products such as aldehydes, sulfur compounds, and organic acids can generate oxidative stress and favor tumor progression [2,3,12–15]. Characterizing and comparing the oral bacterial metagenome in patients with OSCC is essential for understanding the role of the microbiome in oral oncogenesis. Although various bacterial genera and species associated with oral cancer have been identified, further research is needed to determine the function of bacterial genes and products and their interaction with the tumor microenvironment. Implementing functional studies will allow the identification of new therapeutic targets and lead to the development of more effective prevention strategies and treatments [16–18]. Alterations in the composition and structure of the oral microbiome have been associated with the development of oral cancer. In the 2017 study by Wolf et al. [19], a significant increase in the abundance of the bacterial genera *Actinomyces, Schwartzia, Treponema*, and *Selenomonas* was observed in the saliva of patients with OSCC compared to healthy individuals. These findings suggest that these bacterial genera could serve as potential biomarkers for the early diagnosis of OSCC and as possible therapeutic targets. While Singh et al. in 2023 [20] evaluated the dynamics of the oral microbiome at different stages of oral cancer, from precancerous lesions to advanced stages, and its impact on the tumor immune response. The researchers analyzed the bacterial composition of 95 tissue samples using 16S rRNA sequencing. The results showed an enrichment of the genera *Streptococcus* and *Rothia* in precancerous lesions, while *Capnocytophaga, Fusobacterium*, and *Treponema* were more abundant in tumors. Additionally, a higher presence of *Capnocytophaga* was observed in advanced stages and *Fusobacterium* in early stages. Immunohistochemical analyses suggested that the tumor microbiota induces an immunosuppressive microenvironment, promoting cancer progression.

Therefore, the characterization of the bacterial microbiome in OSCC using rapid and efficient technologies will allow us to understand the leading role of bacterial species in complex diseases such as cancer. In this study, the bacterial microbiome of plaque, saliva, and tumor tissue was characterized by using metagenomic sequencing in patients with OSCC and dental plaque and saliva in patients without OSCC.

Oral metagenomics provide a broader framework for understanding microbial involvement in carcinogenesis because it enables the simultaneous characterization of taxonomic composition and functional potential. Oral metagenomic profiling can identify microbial signatures associated with cancer prediction, diagnosis, and personalized therapeutic strategies. Shotgun metagenomics offers higher taxonomic resolution, improved species level detection, and direct identification of genes involved in virulence, metabolism, DNA repair, oxidative stress responses, and host microbe interactions than 16S rRNA amplicon sequencing. This distinction is important because the microbial contribution to oral carcinogenesis may depend less on the mere presence of specific taxa and more on their functional activities within tumor-associated niches. Shotgun metagenomic and metatranscriptomic studies have shown that cancer-associated oral communities may display altered functional potential and increased expression of virulence- and metabolism-related pathways in OSCC, supporting the concept that microbial functional remodeling participates in tumor microenvironment adaptation. Therefore, metagenomic approaches complement amplicon-based surveys by moving beyond taxonomic composition toward microbial functional potential [21–23]. Previous metagenomic studies have identified extensive alterations in the oral bacteriome of patients with OSCC. For example, one study identified 1,269 bacterial OTUs and performed a comparative phylogenetic analysis to evaluate the association of individual taxa with health and disease. The authors reported that numerous bacterial species were associated with eubiosis in healthy controls, whereas patients with OSCC exhibited a higher abundance of dysbiosis-associated taxa. These findings support the concept that oral microbial communities undergo substantial compositional changes during OSCC development and highlight the potential of specific bacterial species as biomarkers of disease and microbial dysbiosis [24].

## METHODS

### Sample Collection

A cross-sectional study was conducted in 10 patients with OSCC (dental plaque, saliva, and tumor tissue) and 10 patients without OSCC (dental plaque and saliva). Patients with OSCC were recruited from Hospital Universitario San Ignacio, the National Cancer Institute in Bogotá, Colombia, and Hospital of Caldas in Manizales, Colombia. Patients without OSCC were recruited from the Faculty of Dentistry at Pontificia Universidad Javeriana in Bogotá, Colombia, in 2016 and 2017. The sample size was determined based on convenience because the study did not aim to establish associations.

The inclusion criteria for patients with cancer were any age and sex, confirmed diagnosis of oral squamous cell carcinoma by histopathological study, a minimum of four natural teeth, and voluntary participation with signed informed consent.

Smoking and alcohol consumption were recorded as clinical and epidemiological variables because of their recognized association with OSCC; however, they were not considered inclusion criteria or prerequisites for patient recruitment. The exclusion criteria were as follows: patients who had received treatment for the neoplasm, antibiotic treatment in the last 3 months, or orthodontic or periodontal treatments in the last year and poor-quality DNA sequences. For the control group, participants were selected based on age similarity to OSCC patients, absence of periodontal disease, no history of orthodontic treatment, and no recent antibiotic use. Healthy controls were recruited from the participating institutions during the same study period. Smoking and alcohol consumption are recognized risk factors for OSCC; therefore, these variables were recorded for all participants and considered during control selection to achieve the greatest possible comparability between cases and controls. Sample collection was initiated following approval from the bioethics committee and informed consent from all participants (CIEFOPUJ Act 11 of 2015). Dental plaque, saliva, and tumor tissue samples were obtained from patients with OSCC, whereas only plaque and saliva samples were collected from control individuals. All samples were collected over a 12-month period using a standard protocol. Participants provided approximately 5 mL of unstimulated saliva, which was collected in tubes containing RNA later buffer to preserve nucleic acids. Each participant completed a dental chart (odontogram) to record the presence or absence of teeth. Participants were instructed not to brush their teeth the night before sample collection to ensure accurate assessment of dental plaque. The modified Silness and Löe Index was used to evaluate plaque accumulation. Supragingival plaque was collected using a scraping technique with Gracey Micro Mini Five curettes, with cotton rolls and a triple syringe used for isolation and drying. The plaque was sampled from the buccal, distal, palatal/lingual, and distal surfaces of teeth 16, 21, 24, 36, 41, and 44. The mesial surface of an adjacent tooth was sampled if any of these teeth were missing. Individuals with only four teeth were designated as index teeth. All plaque samples were preserved in RNA later buffer and stored at −70 °C until further analysis. Among the limitations of this study, potential confounding variables, including host immune factors, sex, oral hygiene, dental and periodontal status, dietary habits, smoking, and alcohol consumption, were considered.

### DNA extraction, quantification, and analysis of quality

Genomic DNA was extracted from approximately 1–5 mg of fresh tumor tissue using the MasterPure™ Complete DNA and RNA Purification Kit (epicenter Biotechnologies®, Madison, WI, USA), following the manufacturer’s instructions with a minor modification to improve bacterial cell lysis. Tissue fragments were suspended in 300 μL of Tissue and Cell Lysis Solution and mechanically homogenized using a filtration-based homogenization system. The homogenate was subjected to a preliminary lysozyme digestion before the standard lysis procedure to enhance the disruption of bacterial cell walls.

Subsequently, 2 μL of Proteinase K (50 μg/μL) were added to digest the structural and nucleoproteins, thereby promoting complete lysis of both the host and bacterial cells. The samples were incubated at 65 °C for 30 min, with gentle mixing every 5 min, according to the manufacturer’s instructions. Cellular proteins and debris were removed using MPC Protein Precipitation Reagent, and genomic DNA was recovered by isopropanol precipitation, washed with 70% ethanol, air dried, and resuspended in TE buffer. DNA concentration and purity were determined using spectrophotometric analysis (Qubit Fluorometer, Thermo Fisher Scientific), and DNA integrity was verified before library preparation and sequencing. DNA samples that met the quality criteria were subjected to high-throughput metagenomic sequencing using the Illumina MiSeq platform. The Illumina Nextera XT DNA Library Preparation Kit (Illumina, San Diego, CA, USA) was used to prepare DNA libraries. Libraries with an average fragment size of approximately 250 bp were generated by enzymatic tagmentation and amplified according to the manufacturer’s recommendations using the standard 12-cycle indexing PCR. The libraries were quantified, normalized, pooled at equimolar concentrations, and sequenced on an Illumina MiSeq™ platform. We evaluated raw sequencing reads using FastQC to assess sequence quality, read length, GC content, and adapter contamination before downstream bioinformatic analyses. Negative extraction controls and mock microbial community standards were not included in the library preparation or sequencing. Briefly, the DNA was fragmented, adapters were added, and PCR was used to generate sequencing libraries. All libraries were quantified and pooled in equimolar concentrations. Automated cluster generation and paired-end sequencing were performed on the Illumina MiSeq™ platform according to the manufacturer’s instructions.

### Bioinformatic analysis of the metagenome

Raw sequencing data in FASTQ format were processed using a standardized bioinformatics workflow implemented in fastp for adapter removal and quality filtering. Reads with a Phred quality score below 20 were trimmed, low-quality bases at the 3′ ends were removed, and reads shorter than 100 bp were discarded after trimming. The sequence quality before and after preprocessing was assessed using FastQC, and the quality reports were summarized using MultiQC. Reads assigned to Homo sapiens were removed before taxonomic profiling to minimize host contamination. We performed taxonomic classification using Kraken2 v2.1.3 with the PlusPF database (k2_pluspf_20260226), and refined species-level abundance estimates using Bracken v2.9. To reduce potential sequencing artifacts, taxa represented by fewer than 10 reads or detected in only a single sample were excluded before downstream analyses. The complete bioinformatics workflow, including software versions and analytical scripts, is publicly available in the project’s GitHub repository [25].

### Diversity Analysis

Beta diversity was assessed using Bray–Curtis dissimilarity followed by principal coordinate analysis (PCoA) to visualize patterns of microbial community composition among sample groups. PCoA was used solely as an exploratory visualization because of the modest sample size (n = 10 per group) and the primary objective of identifying differentially abundant taxa rather than testing global differences in community structure.

### Statistical Analysis

Relative bacterial abundances were calculated from the RF feature table. Differences in bacterial abundance between patients with OSCC and healthy controls were evaluated using the Mann–Whitney U test, and p-values were adjusted for multiple testing using the Benjamini–Hochberg false discovery rate (FDR) procedure. Only taxa that remained significant after FDR correction (q < 0.05) were considered differentially abundant, whereas taxa that did not meet this threshold were interpreted as descriptive trends. PCoA based on Bray–Curtis dissimilarity was used to visualize differences in microbial community composition among sample groups, while data processing, diversity analyses, and graphical visualizations were conducted in R using the phyloseq package and complementary packages for statistical analysis and visualization. The microbial profiles of patients with and without OSCC were summarized using descriptive statistics, including means, standard deviations, and relative abundance distributions.

### Limitations

Differences in oral hygiene between patients and controls may have acted as a confounding factor influencing the observed microbial profiles. Because of the observational cross-sectional design and limited sample size, the adjustment for potential confounders through multivariable analyses was not feasible. The predominance of female participants (90%) may limit the external validity and generalizability of the findings.

## RESULTS

### Validation and quality control of data

The average reads initially obtained from patients with OSCC were 97.746,6 bp in dental plaque, 99.045,6 bp in saliva, and 76.752 bp in tumor. After data cleaning, the average reads obtained were 96.957 bp in dental plaque, 97.763,2 bp in saliva, and 75.294,2 bp in tumor, with a mean length of 312,33 bp. In patients without OSCC, the initial average reads were 100.275.8 bp in dental plaque and 94.391.88 bp in saliva. After data cleaning, the average reads obtained were 99.683.38 bp in dental plaque and 93.623.13 bp in saliva, with a mean length of 311.5 bp (Table 1). The complete bioinformatics workflow, including software versions, databases, parameters, and analysis scripts, is publicly available at the project’s GitHub repository [25]. A total of 49 biological samples were initially processed.

**Table 1.** Number of reads before and after the bioinformatic analysis.

| Group | Number of initial reads |  |  |  | Bioinformatic parameters | Number of final reads |  |  |  |
| --- | --- | --- | --- | --- | --- | --- | --- | --- | --- |
|  | Plaque | Saliva | Tumor | Length pb |  | Plaque | Saliva | Tumor | Length pb |
| **Patients (Mean)** | 97746,6 | 99045,6 | 76752 | 300 | Fastp: Q≥30; W=4; L=50 bp<br>Kraken2: conf. = 0.1<br>Bracken: r = 50; S; t = 10 | 96957 | 97763,2 | 75294,2 | 100-266 |
| **Controls (Mean)** | 100275,8 | 94391,88 | NA | 300 |  | 99638,3 | 93623,1 | NA | 100-242 |

During sequence quality assessment, three saliva samples from healthy controls failed to meet the predefined sequencing quality criteria required for reliable downstream metagenomic analyses and were therefore excluded before taxonomic profiling and statistical analyses. Consequently, the final analytical dataset consisted of 46 high-quality samples, all of which were deposited in the National Center for Biotechnology Information (NCBI) BioProject PRJNA1165312.

The study population primarily comprised women (90%), with a mean age of 56 years. Notable disparities in oral hygiene practices were observed: 80% of the control group and 20% of the patient group reported good oral hygiene. In contrast, 60% of the patients exhibited fair to poor oral hygiene. Regarding tumor localization, 50% were located on the tongue, 30% on the cheek, and 20% on the lips (Table 2). Not all individuals retained the six index teeth (16, 21, 24, 36, 41, and 44) due to the participants’ age and previous tooth loss. Among patients with OSCC, 2 of the 10 participants (20%) retained all six index teeth, 2 (20%) retained five index teeth, 1 (10%) retained four index teeth, and 5 (50%) retained two index teeth. In the control group, 2 of the 10 participants (20%) retained all six index teeth, 4 (40%) retained five index teeth, and 4 (40%) retained four index teeth. All participants had at least two available index teeth for assessment; therefore, no individual was excluded from the study because of missing index teeth. Plaque sampling was successfully completed in all participants using the available index teeth according to the predefined sampling protocol.

**Table 2.**
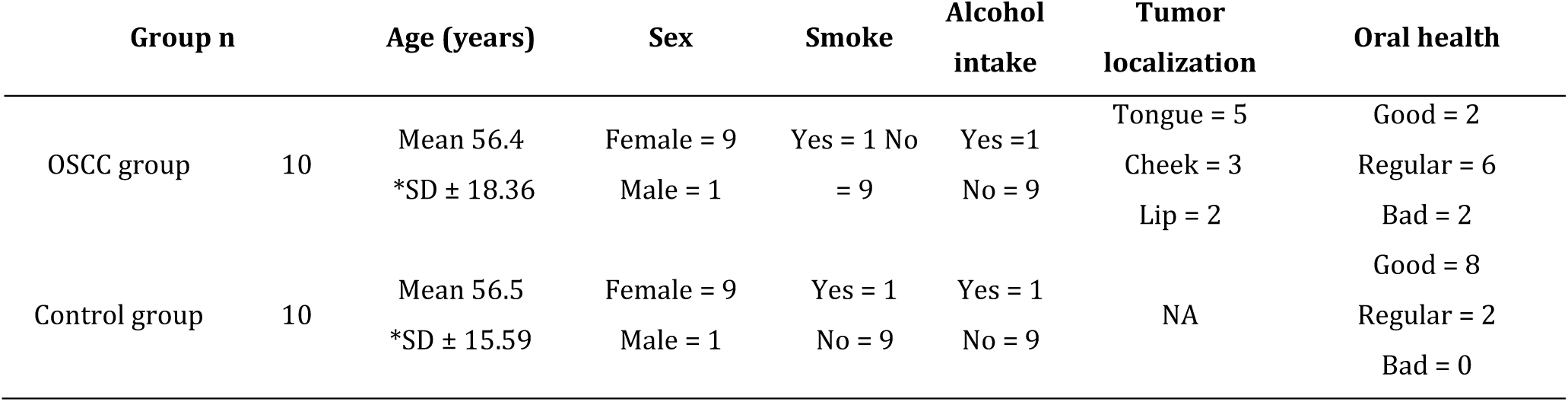
Clinical characteristics of patients with and without OSCC.

Because oral hygiene differs substantially between patients and controls, the observed microbial differences cannot be attributed exclusively to OSCC. Poor oral hygiene influences oral microbial composition and may have contributed to the microbial profiles identified in this study. Adjustment for this potential confounding factor was not feasible given the limited sample size. Therefore, the findings should be interpreted as exploratory and should be confirmed in larger, well-matched cohorts.

### Bacterial Metagenome

The FastQC analysis indicated that most reads in the metagenomes exhibited high Phred scores (≈28–32) across the majority of their length, reflecting overall good sequencing quality. However, a progressive decline in quality was observed toward the 3′ ends of the reads, particularly after ∼380–400 bp, where the Phred scores pronouncedly dropped and the variability among samples increased. In contrast, the initial bases generally maintain high-quality values, with no evidence of systematic reduction (Figure 1a). The GC content distribution of the metagenomes shows that most sequences cluster within a relatively narrow range of approximately 48%–60% GC, with prominent peaks centered around ∼52–56% across samples. The distributions are generally unimodal and largely overlap, indicating a consistent overall GC profile among the datasets. Although minor shifts in peak position and height were observed between samples, there was no evidence of extreme outliers or secondary peaks that would clearly suggest contamination was observed. These subtle variations likely reflect differences in metagenomic composition among samples rather than technical artifacts (Figure 1b). The FastQC graph for mean sequence quality scores in the metagenomes shows that most sequences exhibit high average quality, with Phred scores clustering predominantly between 28 and 31. The distributions peak around ∼29–30, indicating consistently good overall quality across the samples. A small proportion of the sequences fell within the moderate quality range (20–28), and virtually none were observed in the low-quality range (<20). Overall, these results indicate that the datasets have high mean sequence quality, suggesting that minimal filtering must ensure reliable downstream analyses (Figure 1c). The graph of the read length distribution (FastQC) shows the distribution of sequence lengths (bp) versus read count across the 49 samples analyzed. A bimodal distribution is observed, with a predominant peak between ∼50–90 bp and a second group of reads between ∼340–370 bp. The high abundance of short reads suggests the presence of truncated fragments or processing-derived products, whereas the longer reads likely correspond to fragments of the expected library length (Figure 1d).

**Figure 1.**
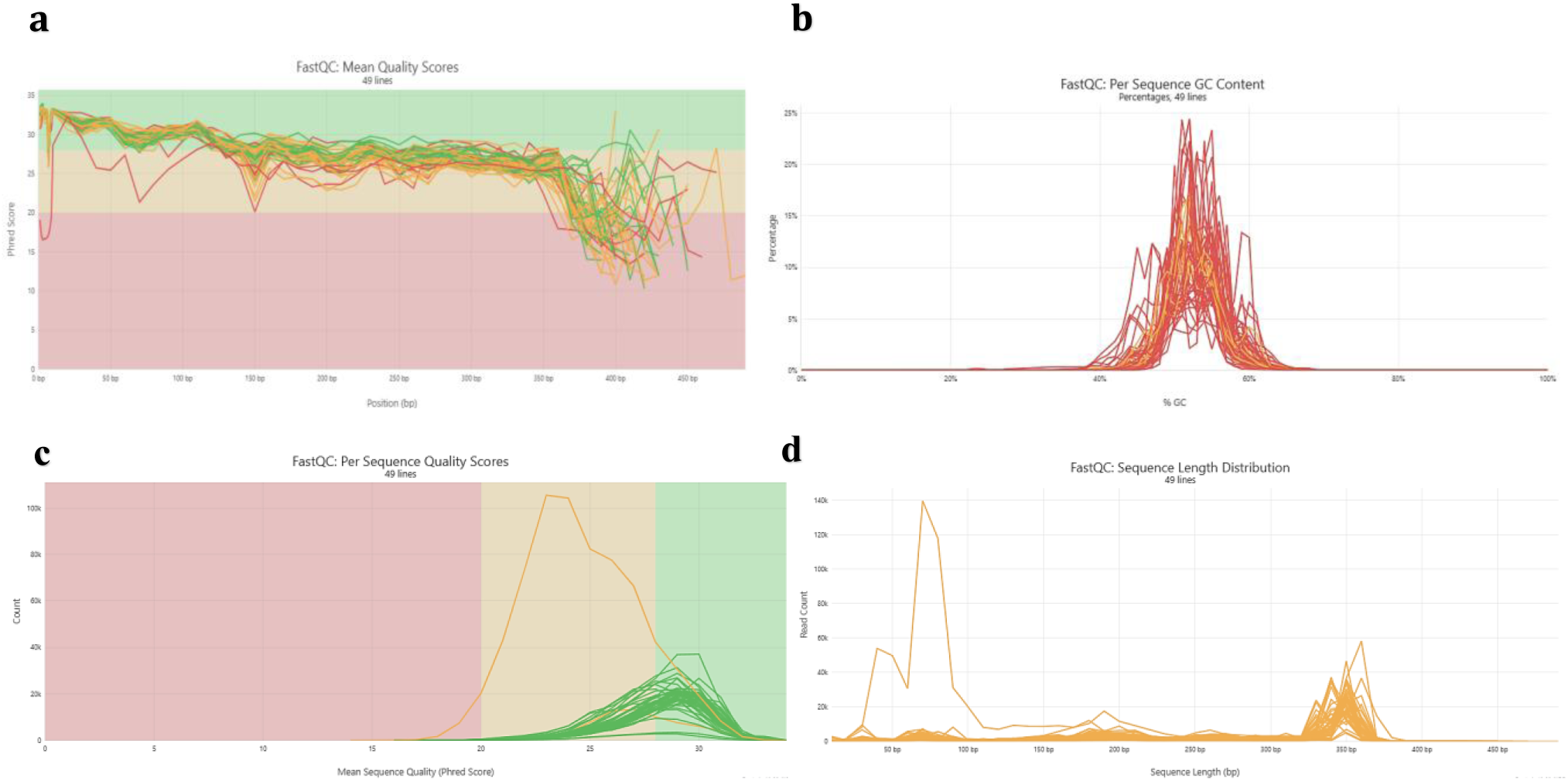
Analysis of metagenome data

Comparisons of the number of bacterial species found in metagenomes from dental plaque, saliva, and tumors in patients with OSCC and dental plaque and saliva in patients without OSCC revealed differences in bacterial composition. In patients with OSCC, 164 bacterial species were identified. Of these, all three sample types contained 65 species (Figure 2a, purple).

**Figure 2.**
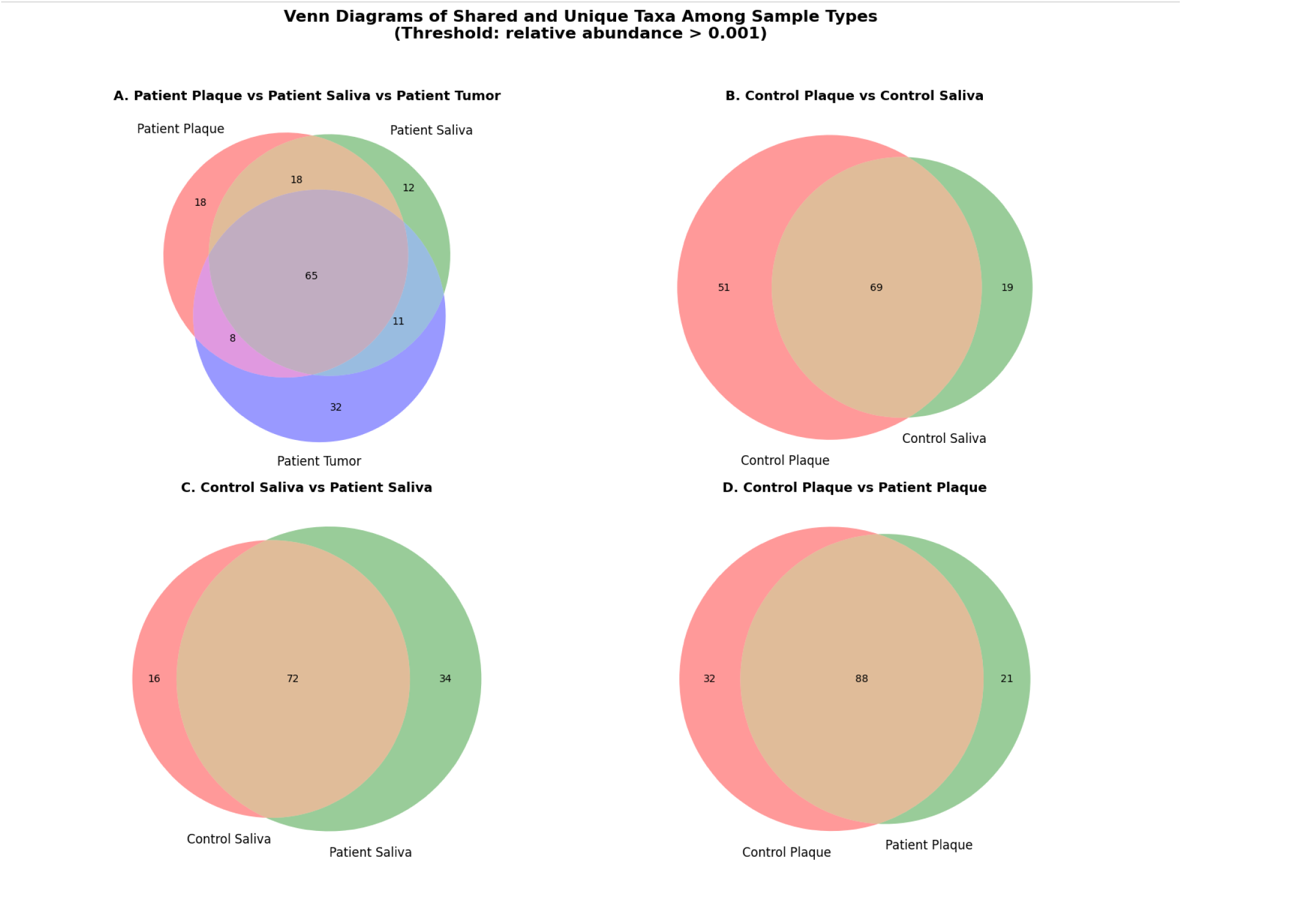
Bacterial species present in dental plaque, saliva, and tumor samples from patients with and without OSCC.

Comparisons of the number of bacterial species found in metagenomes from dental plaque, saliva, and tumors in patients with OSCC and dental plaque and saliva in patients without OSCC revealed differences in bacterial composition. In patients with OSCC, 164 bacterial species were identified. Of these, all three sample types contained 65 species (Figure 2a, purple). Additionally, 18 species were exclusive to dental plaque (Figure 2a, pink), 12 to saliva (Figure 2a, green), and 32 to tumors (Figure 2a, lilac). Furthermore, 18 species were shared between dental plaque and saliva, 8 between dental plaque and tumors, and 11 between saliva and tumors (Figure 2a). Comparative analyses of dental plaque and saliva metagenomes in patients without OSCC identified 139 bacterial species. Of these, 69 species were shared between both sample types (Figure 2b, brown), while 51 were exclusive to dental plaque (Figure 2b, pink) and 19 to saliva (Figure 2b, green). Comparisons between saliva metagenomes from OSCC and non-OSCC patients revealed 122 shared species (Figure 2c, brown), with 34 species more frequent in OSCC saliva (Figure 2c, green) and 16 in non-OSCC saliva (Figure 2c, pink). Similarly, comparisons of dental plaque metagenomes between patients with and without OSCC showed 141 shared species (Figure 2d, brown), while 21 species were more abundant in patients with OSCC plaque (Figure 2d, green) and 32 in patients with non-OSCC plaque (Figure 2d, pink). Bacterial species present in plaque, saliva, and tumor samples from patients were compared with those found in plaque and saliva samples from control individuals. The most representative species in dental plaque and saliva from patients with OSCC were *Schaalia sp. HMT-172, Leptotrichia sp. oral taxon 498*, and *Pseudoleptotrichia goodfellowii*, which showed relatively higher CLR-transformed abundances in these sample types. In contrast, *Streptococcus cristatus* exhibited higher abundance in plaque samples from control individuals.

When comparing bacterial species present in the dental plaque of patients with and without OSCC, taxa such as *Pauljensenia hongkongensis*, *Capnocytophaga sp. oral taxon 878*, and *Streptococcus mutans* were notable, particularly in patient plaque samples.

Additionally, *Cardiobacterium hominis* and *Treponema sp. Marseille-Q4132* displayed variable abundances across plaque and saliva samples, suggesting differences in microbial composition between groups. Notably, the tumor samples were characterized by an increased abundance of species such as *Escherichia coli, Lacrimispora sp. HJ-01*, and *Sellimonas catena*, indicating a distinct microbial profile compared with the oral samples (Figure 3). Overall, these patterns indicate that microbial composition differs between OSCC and control samples, with specific taxa enriched in patient plaque and tumor environments, while others are more characteristic of control-associated oral niches (Figure 3).

**Figure 3.**
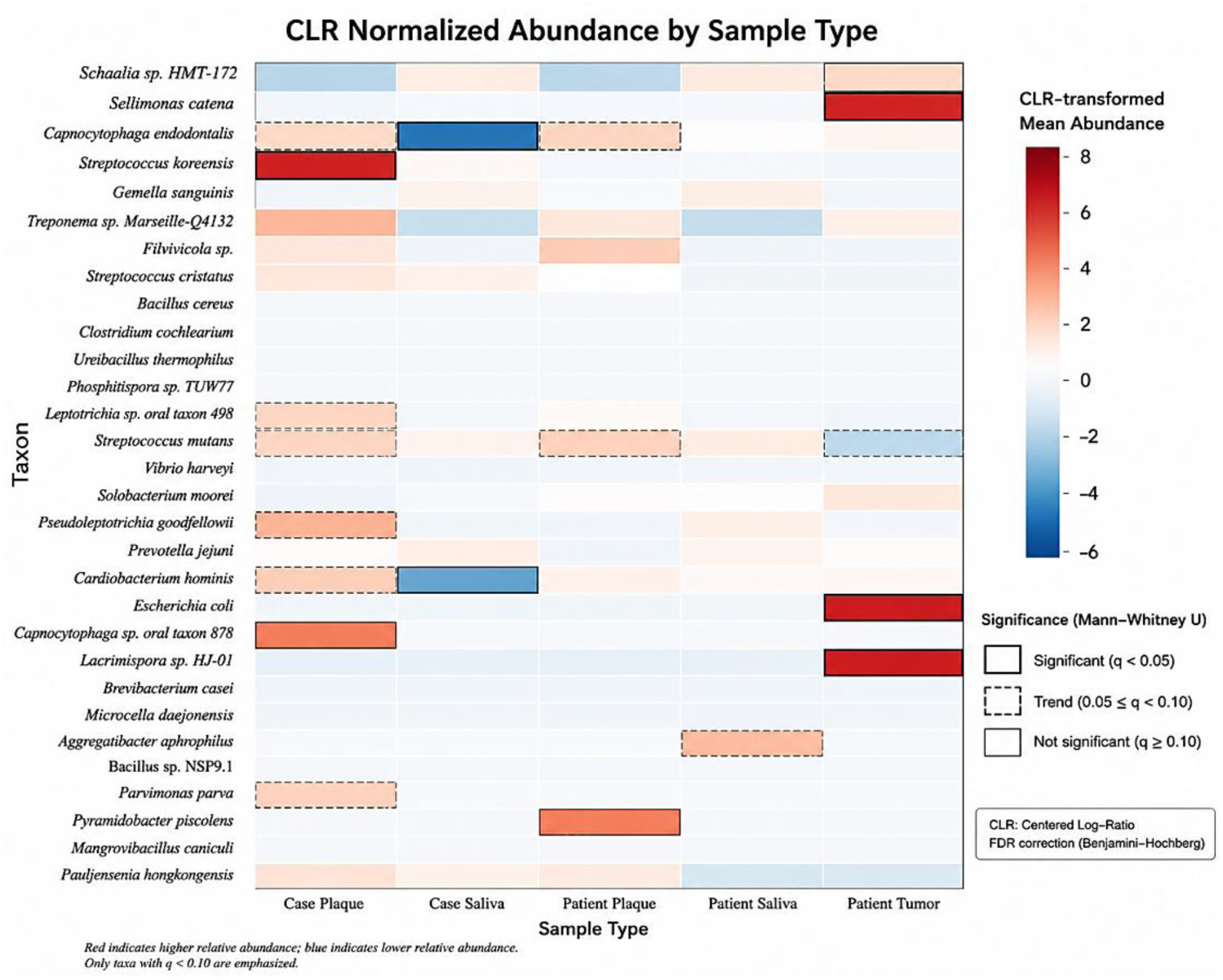
Bacterial species diversity in saliva, dental plaque, and tumor samples from patients with and without OSCC

### Predominant bacterial taxa distribution according to oral microenvironment

The CLR-normalized heatmap revealed clear differences in bacterial composition among dental plaque, saliva, and tumor tissue, indicating that each oral microenvironment harbored a characteristic bacterial community. Although these findings are exploratory, several taxa exhibited preferential abundance in specific niches.

Dental plaque exhibited the highest taxonomic diversity and the highest number of taxa with elevated relative abundance. The control plaque was characterized by the predominance of *Streptococcus koreensis*, which showed the highest abundance among all plaque-associated taxa. Other highly abundant species included *Capnocytophaga sp*. oral taxon 878, *Treponema sp*. *Marseille* Q4132, *Pseudoleptotrichia goodfellowii*, *Cardiobacterium hominis*, *Leptotrichia sp*. oral taxon 498, *Parvimonas parva*, *Streptococcus mutans*, and *Pauljensenia hongkongensis*. Taxa such as *Gemella sanguinis, Streptococcus cristatus, Prevotella jejuni, Fluviicola sp., Capnocytophaga endodontalis,* and *Schaalia sp*. HMT-172 were found at moderate abundance, whereas virtually no tumor-associated taxa were found, including *Sellimonas catena*, *Escherichia coli*, and *Lacrimispora sp*. HJ-01. In contrast, the patient plaque displayed a partially distinct bacterial profile. *Pyramidobacter piscolens* was the most abundant taxon in this microenvironment, followed by *Fluviicola sp., Parvimonas parva, Pauljensenia hongkongensis, Capnocytophaga endodontalis, Treponema sp. Marseille* Q4132, and *Schaalia sp.* HMT-172.

*Gemella sanguinis* and *Streptococcus mutans* remained moderately abundant, whereas several taxa that predominated in the control plaque, including *Streptococcus koreensis, Cardiobacterium hominis, Pseudoleptotrichia goodfellowii,* and *Capnocytophaga sp.* oral taxon 878, showed comparatively lower abundance. Overall, the plaque samples were dominated by bacteria associated with mature oral biofilms and anaerobic periodontal communities (Figure 3). In control saliva, the predominant taxon was *Capnocytophaga endodontalis*, which reached its maximum abundance in this sample type. Other relatively abundant microorganisms included *P. jejuni, G. sanguinis, P. hongkongensis, S. mutans,* and *S. cristatus*. In contrast, plaque-associated taxa, such as *Streptococcus koreensis, Treponema sp. Marseille* Q4132, Cardiobacterium hominis, and *Capnocytophaga sp. oral taxon* 878, were less abundant. Similarly, patient saliva exhibited a relatively uniform taxonomic profile without a single dominant species. The most abundant taxa included *Aggregatibacter aphrophilus, Pseudoleptotrichia goodfellowii, Prevotella jejuni, Streptococcus mutans, Gemella sanguinis*, and *Capnocytophaga endodontalis,* whereas *Schaalia sp.* HMT-172 and *Cardiobacterium hominis* were detected at intermediate abundance.

Most plaque-associated anaerobes were less abundant than dental plaque, suggesting that saliva supports a more evenly distributed bacterial community (Figure 3). Tumor tissue exhibited the most distinctive bacterial composition among the oral microenvironments analyzed. The bacterial community was dominated by S. *catena,* which showed the highest relative abundance in tumor samples. Additional taxa exhibiting marked abundance included *Escherichia coli, Lacrimispora sp. HJ-01, Capnocytophaga sp.* oral taxon 878, and *Solobacterium moorei*, all of which reached their maximum abundance in tumor tissue (Figure 3). The moderately abundant taxa included *Schaalia sp*. HMT-172, *Treponema sp*. *Marseille* Q4132, *Gemella sanguinis,* and *Capnocytophaga endodontalis.* Conversely, microorganisms that predominated in dental plaque, including *Streptococcus koreensis, Leptotrichia sp.* oral taxon 498, *Cardiobacterium hominis, Pseudoleptotrichia goodfellowii, Parvimonas parva, Pyramidobacter piscolens,* and *Pauljensenia hongkongensis,* exhibited comparatively low abundance in tumor tissue.

Taken together, these exploratory analyses indicate that dental plaque harbors the greatest bacterial diversity and the largest number of abundant taxa, particularly species commonly associated with mature oral biofilms and periodontal communities. Salivary samples exhibited a more homogeneous microbial composition, with relatively few dominant taxa and lower overall heterogeneity. In contrast, tumor tissue was characterized by a distinct bacterial profile dominated by a limited number of taxa, including *Sellimonas catena*, *Escherichia coli*, *Lacrimispora* sp. HJ-01, *Capnocytophaga* sp. oral taxon 878, and *Solobacterium moorei*, highlighting the existence of different bacterial community profiles.

Distribution of key alpha diversity metrics across the 49 metagenomic samples. Shannon diversity (top-left): Most samples have Shannon values between ∼1.5 and 3.0, with a median of approximately 2.5. This indicates moderate diversity. Some samples (e.g., 26CP, N19CP) show higher diversity (>3.0), whereas others (e.g., 5MS, 31MT) have very low diversity (<1.5), suggesting that few taxa dominate. Observed features/richness (top-right): The number of detected species ranges widely from ∼10 to nearly 200. The median is approximately 50 species per sample. The unknown_barcode sample is a clear outlier with ∼193 species, indicating much higher richness than the rest of the dataset. Chao1 Richness Estimator (bottom-left): Very similar to the observed richness. Chao1 accounts for rare species, so the fact that it is close to the observed features in most samples suggests that rare taxa are not heavily underestimated. Berger-Parker Dominance (bottom-right): Most samples have dominance values between 0.2 and 0.4 (median ∼0.28), indicating that one taxon typically represents 20%–40% of the community. Higher values (closer to 0.7 in 5MS) indicate strong dominance by a single species (likely Streptococcus or similar in your case) (Figure 4). The Bray–Curtis principal coordinates analysis (PCoA) provided an exploratory visualization of the composition of the microbial community across sample types. Differences in sample distribution were visually apparent among oral microenvironments; however, these patterns should be interpreted descriptively rather than as evidence of statistically significant group separation because no formal multivariate significance test was performed (Figure 4B).

**Figure 4.**
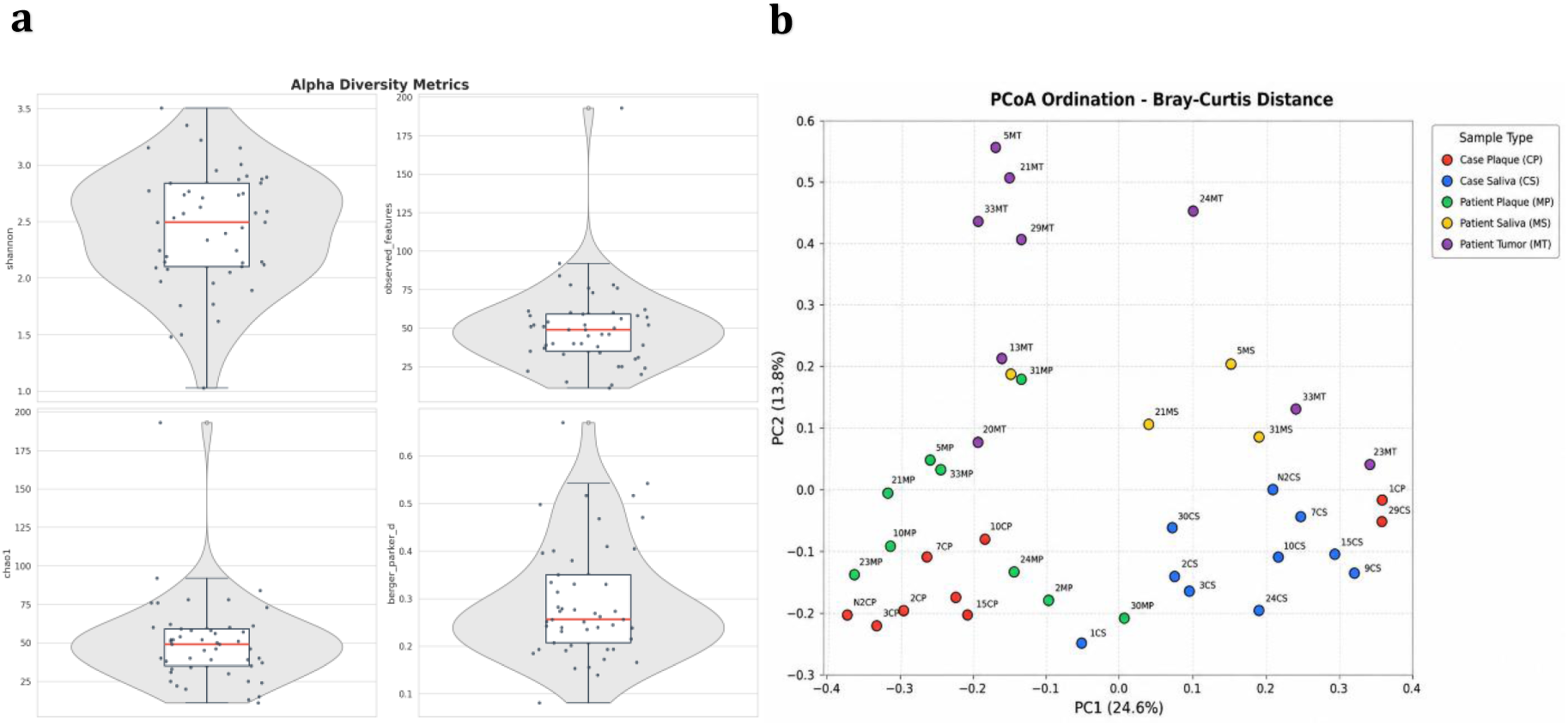
**a**. Alpha diversity metrics of bacterial species in saliva, dental plaque, and tumor samples from patients with and without OSCC. **b.** Exploratory Bray–Curtis Principal Coordinate Analysis showing similarities among microbial communities. No statistical group comparison was performed.

## DISCUSSION

Oral biodiversity is estimated to encompass over 700 bacterial species. Both mutualistic and pathogenic bacteria can coexist within complex biofilm communities, contributing to homeostasis maintenance [3, 26]. Most bacteria inhabit these biofilms, which dominate the oral cavity’s ecological niches. Because oral hygiene differed substantially between patients and controls, the observed microbial differences cannot be attributed exclusively to OSCC.

Poor oral hygiene influences oral microbial composition and may have contributed to the microbial profiles identified in this study. Adjustment for this potential confounding factor was not feasible given the limited sample size. Therefore, the findings should be interpreted as exploratory and should be confirmed in larger, well-matched cohorts. Host-related factors may disrupt the delicate balance within these niches, potentially leading to ecological imbalances in the bacterial community and resulting in oral disease. Such imbalances, known as dysbiosis, favor the growth of pathogens that generate inflammatory conditions capable of damaging tissue cells through their virulence and pathogenicity factors [27]. Chronic inflammation, a hallmark of cancer, can be promoted by an altered oral flora, driving mutagenesis, angiogenesis, and uncontrolled cell proliferation, ultimately contributing to oral cancer development [27, 28]. Furthermore, these imbalances may induce epigenetic alterations, cellular degeneration, and malignant transformation.

The oral microbiota is increasingly recognized as a key factor in the development and progression of OSCC, and dysbiosis has been associated with inflammation and tumorigenesis, highlighting the potential of microbial alterations as tools for early, noninvasive diagnosis [29, 30]. The identification of bacteria with oncogenic potential, such as *Fusobacterium nucleatum* and *Porphyromonas gingivalis*, which have been associated with OSCC, may contribute to a better understanding of the mechanisms by which microbial communities participate in tumor initiation and progression, including the promotion of chronic inflammation, modulation of host immune responses, alteration of cell signaling pathways, and inhibition of apoptosis [31].

To the best of our knowledge, compared with previous studies, which have primarily evaluated individual oral sample types, our study simultaneously characterized bacterial communities in dental plaque, saliva, and tumor tissue within the same cohort. This approach revealed differences in bacterial community profiles among oral microenvironments, demonstrating that several taxa exhibited preferential abundance according to the oral niche rather than being uniformly associated with OSCC. These findings expand the current knowledge of the ecological organization of the oral microbiome in OSCC [32]. The potential involvement of taxa such as *Sellimonas catena*, *Lacrimispora* sp. HJ-01, *Pyramidobacter piscolens*, and *P. hongkongensis* in OSCC remains poorly characterized. Therefore, the preferential distribution across specific oral microenvironments observed in this study should therefore be considered exploratory and warrants validation in larger cohorts [32].

Our findings are partially consistent with those reported by Fukase et al. [33]. who demonstrated significant alterations in the oral microbiome of patients with OSCC and observed enrichment of *Treponema* spp. in patients with advanced disease.

Similarly, our study identified relatively higher abundance of periodontal-associated taxa, including *Treponema* spp., *Parvimonas parva*, and *Capnocytophaga endodontalis*, in dental plaque samples from patients with OSCC. While the enrichment of *Treponema* is consistent with previous observations, the increased abundance of *P. parva* and *C. endodontalis* in our cohort extends these findings and supports the potential involvement of periodontal-associated bacterial communities in the oral tumor microenvironment. In contrast, the preferential distribution of taxa such as *Sellimonas catena*, *Lacrimispora* sp. HJ-01, *Pyramidobacter piscolens*, and *Pauljensenia hongkongensis* across distinct oral microenvironments represents an exploratory observation that has not been extensively characterized in the current literature on OSCC. These findings highlight the importance of considering multiple oral niches when investigating OSCC-associated microbiome alterations [33]. In this study, metagenomic analyses revealed distinct bacterial communities across the oral microenvironment. The dental plaque from patients with OSCC was characterized by a relatively higher abundance of *Pyramidobacter piscolens*, *Fluviicola* sp., *Parvimonas parva*, *Pauljensenia hongkongensis*, and *Capnocytophaga endodontalis*, whereas the control plaque was enriched in *Streptococcus koreensis*, *Cardiobacterium hominis*, *Pseudoleptotrichia goodfellowii*, and *Capnocytophaga* sp. oral taxon 878. The tumor tissue exhibited a distinct bacterial profile dominated by *Sellimonas catena*, *Escherichia coli*, *Lacrimispora* sp. HJ-01, and *Solobacterium moorei*, supporting the existence of different bacterial community profiles in OSCC.

This study should be interpreted in light of several limitations. First, the relatively small number of participants limited the statistical power to detect modest differences in bacterial abundance and precluded adjustment for potential confounding variables as an exploratory pilot study based on a convenience sample. Consequently, the reported microbial associations should be hypothesized and validated in larger, well-matched, multicenter cohorts. Second, although all samples underwent rigorous quality control and standardized bioinformatic processing, the sequencing depth may have reduced the detection of low-abundance taxa, particularly in tumor samples. In addition, the absence of negative extraction controls and mock microbial community standards prevented the direct evaluation of potential background contamination. Finally, differences in oral hygiene between patients and controls may have contributed to the observed microbial profiles independent of OSCC status. Similarly, the predominance of female participants limits the findings’ external validity. These factors should be considered when interpreting the ecological differences observed among the oral microenvironments.

### Re-use potential

These data provide species-level taxonomic profiles that can be integrated with other metagenomic and metatranscriptomic datasets to investigate the role of the oral microbiome in the pathogenesis of OSCC. The complete bioinformatics workflow is fully reproducible, with all software versions, databases, and analytical procedures described in the Methods section. The Conda-based pipeline and analysis scripts are publicly available in the metagen GitHub repository [25], facilitating data reuse, independent validation, and future comparative studies.

## Data Availability

The metagenomic sequencing data generated in this study are publicly available in the National Center for Biotechnology Information (NCBI) BioProject database under accession number PRJNA1165312 (Project ID: 1165312) [34].

**Availability of Source Code and Requirements.** https://github.com/aidbiolab/metagen

## Ethics and Informed Consent

The Ethics Committee of the Pontificia Universidad Javeriana, Bogotá D.C., Colombia, approved this study (approval number OD-0149). All participants signed an informed consent form before the commencement of the study, which clearly and concisely explained the objectives, procedures, possible risks, and benefits associated with their participation.

## Consent for publication

Written informed consent was obtained from all participants.

## Author Contributions

**Conceptualization:** Alveiro Erira, Fredy Gamboa, Adriana García. **Data curation:** Alveiro Erira, Fabian Tobar, José Noguera. **Formal analysis:** Alveiro Erira, Fabian Tobar, José Noguera. **Investigation:** Alveiro Erira, Adriana García, Andrés Ignacio Chala, Andrei Moreno, Eliana Muñoz. **Methodology:** Alveiro Erira, Adriana García, Fredy Gamboa, Fabian Tobar. **Resources:** Alveiro Erira, Adriana García, Andrés Ignacio Chala, Andrei Moreno, Eliana Muñoz, **Software:** Alveiro Erira, Fabian Tobar, José Noguera. **Supervision:** Adriana García, Fredy Gamboa, Alveiro Erira. **Validation:** Alveiro Erira, Fabian Tobar, Adriana García, Fredy Gamboa, Andrés Ignacio Chala, Andrei Moreno, Eliana Muñoz, Angel Cid Arregui. **Visualization:** Alveiro Erira, Fabian Tobar. **Writing – original draft:** Alveiro Erira, Fredy Gamboa, Adriana García. **Writing – review & editing:** All authors.

## Acknowledgements.

The authors would like to express their gratitude to the Pontificia Universidad Javeriana, Bogotá, the Universidad Cooperativa de Colombia, Bogotá, the participating patients and the clinical staff of the collaborating institutions for their assistance with participant recruitment and sample collection.

## Use of AI-assisted tools

During the preparation of this manuscript, OpenAI ChatGPT (GPT-5.5) and Trinka were used to assist with english language editing, improvement of scientific writing and manuscript organization. AI tools were not used to generate, analyze, or interpret the sequencing data, statistical analyses, or scientific conclusions. All AI-assisted outputs were critically reviewed, verified, and revised by the authors, who assume full responsibility for the accuracy, integrity, and final content of the manuscript.

